# SOX9 regulates endothelial tip cell specification to promote cerebral and neuroretinal vascularization

**DOI:** 10.64898/2026.06.07.730751

**Authors:** Typhaine Anquetil, Gael Cagnone, Joel P Howard, Lily Kennepohl, Sahily Rodriguez-Torres, Bruno Larrivée, Alexandre Dubrac

## Abstract

**Background:** Tissue vascularization relies on the organotypic specification of endothelial tip cells to interpret local cues and guide angiogenic sprouts into specific tissue compartments. However, the molecular regulators controlling brain and retinal endothelial tip cell identity remain poorly understood.

**Methods:** Endothelial-specific *Sox9* loss-of-function mouse was combined with single-cell, spatial, and bulk RNA sequencing analyses of developing mouse brain and retinal vasculature, as well as experimental ischemic stroke. Transcriptomic findings were validated using *in situ* hybridization, immunofluorescence, and functional angiogenesis assays.

**Results:** Transcription factor activity analysis of single-cell RNA-sequencing datasets identified SOX9 as a candidate regulator selectively enriched in developing brain and retinal endothelial tip cells. Endothelial-specific deletion of *Sox9* impaired brain and neuroretina vascularization and disrupted the tip cell transcriptomic program, resulting in reduced sprouting angiogenesis and matrix-remodeling pathways. Conversely, SOX9 overexpression in HUVECs promoted neuro-tip-like signatures and enhanced endothelial invasion and sprouting. Following ischemic stroke, single-cell and spatial transcriptomic analyses identified a transient angiogenic endothelial population within the ischemic area. However, these cells failed to express *Sox9* and lacked key developmental brain tip cell features.

**Conclusions:** SOX9 is a key regulator of endothelial tip cell identity and neuronal angiogenesis. These findings reveal fundamental differences between developmental and injury-induced vascular responses and identify SOX9 as a potential therapeutic target to promote functional vascular regeneration.

## INTRODUCTION

Endothelial tip cells drive angiogenic sprouting by sensing environmental cues and guiding nascent vessels ^1–3^. Emerging evidence suggests that tip cells acquire organ-specific properties to ensure that newly formed vessels are functionally adapted to the tissue they perfuse ^4–8^. This concept of organotypic tip cell specialization is particularly relevant in the central nervous system (CNS), where blood vessels also develop barrier properties to protect neuronal homeostasis.

In the brain, angiogenesis is tightly linked to the acquisition of blood-brain barrier (BBB) features, coordinated by signals from the local microenvironment and surrounding cells ^9–11^. During embryogenesis, cerebral vessels sprout from the perineural vascular plexus into the neuroepithelium from E10.5 and progressively acquire BBB properties concomitantly with vascular expansion and neural tissue maturation ^12–14^. After birth, brain vascular development continues extensively to match volumetric brain growth and neuronal circuit maturation. We recently identified three postnatal distinct phases of brain vascular development: an early isometric expansion phase, a regional specialization phase, and a late refinement phase, highlighting that postnatal angiogenesis is tightly coordinated with neurovascular unit maturation and neuronal activity^15,16^.

Beyond development, understanding the mechanisms that govern CNS vascularization is also relevant in pathological context such as ischemic stroke, where angiogenesis and vascular remodeling are suggested to contribute to tissue repair and functional recovery ^17–20^. Although several pro-angiogenic pathways activated after stroke have been identified ^20–22^, the endothelial cell states that emerge during this process remain unclear. Defining the regulators of developmental neurovascular endothelial specialization may therefore provide insight into mechanisms that support vascular regeneration after injury. While canonical pathways such as VEGFA have been extensively studied and shown to regulate sprouting angiogenesis, additional signaling pathways, including WNT and TGFβ, have been implicated in instructing endothelial specialization ^1–4^. We recently demonstrated that neuronal-derived TGFβ signaling controls region-specific postnatal brain angiogenesis ^15^.

In retinal angiogenesis, we also showed TGFβ signaling regulates specialized diving tip cell population (D-tip cells) required for neuroretina deep vascularization. These D-tip cells, which sprout from superficial retinal vessels to invade the neuroretina and form the deep vascular plexus, are transcriptionally distinct from superficial ESM1^+^ tip cells (S-tip cells). D-tip cells display elevated blood-retina barrier gene expression and require endothelial TGFβ/TGFβ receptor I (ALK5) signaling for both their specification and neuroretina vascularization ^4^. Although this work established that tip cell specification is required for proper vascularization of neural tissues, the upstream transcriptional regulator governing this specialized state remained undefined.

Here, we identify the transcription factor SOX9 as a key determinant of neural tip-identity. Using single-cell transcriptomics, genetic loss-of-function, overexpression approach, and functional assays, we show that *Sox9* is selectively enriched in brain- and D-tip cells and is required for their transcriptional specification and angiogenic function. Importantly, *Sox9* is sufficient to reprogram human endothelial cells toward a neuro-tip cell-like state, promoting sprouting and matrix remodeling programs. Interestingly, we found that angiogenic endothelial cells emerging after ischemic stroke fail to express *Sox9* and do not acquire a proper brain-tip cell identity and features amidst pro-angiogenic microenvironment. Together, our findings uncover an organ-specific tip cell identity mechanism and a clear distinction between developmental angiogenesis and injury-induced vascular remodeling.

## Materials and Methods

### Mouse models and experimental protocols

#### Postnatal experiments

All mouse experiments were approved by the Comité Institutionnel des Bonnes Pratiques Animales en Recherche (CIBPAR) of the Azrieli Research Centre of the CHU Sainte-Justine (#2024-6395; #2026-9327). Mice were housed in a pathogen-free facility under a 12 h light/dark cycle at 22 ± 2°C with 40–60% humidity and ad libitum access to food and water. C57Bl/6J (#000664) and *Sox9flox* (#013106) mice were obtained from the Jackson Laboratory. *Sox9flox* mice were crossed with *AplnRCreERT2* mice ^23^ to generate endothelial-specific conditional knockout animals. *Sox9flox;AplnRCreERT2;TdTomato* reporter mice were generated by further crossing with B6.Cg-Gt(ROSA)26Sortm14(CAG-tdTomato)Hze/J reporter mice (Jackson Laboratory, 007914).

To induce postnatal endothelial-specific deletion of *Sox9*, 50 µL of tamoxifen (Sigma, T5648; 2 mg/mL in corn oil) was injected intraperitoneally for three consecutive days and mice were analyzed at the indicated time points. Tamoxifen-injected Cre-negative littermates were used as controls. Unless otherwise specified, both male and female mice were used.

#### Transient Middle Cerebral Artery Occlusion (tMCAO)

Stroke experiments were performed on 8–10-week-old mice of both sexes. Mice received 3 mg/kg buprenorphine 1 h before anesthesia with 2–2.5% isoflurane in a mixture of 70% N2O and 30% O2. After midline neck incision, the internal carotid artery was occluded with a 4-0 nylon monofilament of 18 mm length with a flame-rounded tip (Doccol, 6023910PK10) to occlude the origin of the middle cerebral artery. After 1 h of occlusion, mice were re-anesthetized and the occluding filament was removed to allow for 1, 3, or 14 days of reperfusion ^24^. Successful induction of focal ischemia was confirmed by contralateral hemiparesis. Exclusion criteria were excessive bleeding or death.

#### Tissue collection and preparation

Mice were euthanized at the indicated time points by decapitation (pups) or cervical dislocation (adults). Brains and eyeballs were collected in cold PBS.

For retinal whole-mount immunostaining, eyeballs were fixed in 4% paraformaldehyde (PFA; Fisher Scientific, AAJ19943K2) for 25 min at room temperature (RT), after which retinas were immediately dissected.

For cryosections, brains were fixed in 4% PFA at 4°C overnight, cryoprotected in 30% sucrose (Fisher, BP220-1) for 3 days, embedded in OCT (Fisher Scientific, 23-730-571), and cryosectioned at 30 µm thickness.

### Immunofluorescence

#### Retina whole-mount immunostaining

Retinas were permeabilized and blocked for 1 h at RT in “Claudio” buffer (1% FBS, 3% BSA, 0.5% Triton X-100 (Sigma), 0.01% sodium deoxycholate (Sigma), 0.02% sodium azide in PBS (Sigma)). Primary antibodies were incubated overnight at 4°C and secondary antibodies for 2 h at RT in the same buffer. Retinas were flatmounted using homemade Mowiol mounting medium. Antibodies are listed in Supplementary Table 2.

#### Postnatal brain sections

Floating cryosections (30 µm) were permeabilized and blocked overnight at 4°C in a solution containing 0.5% Triton X-100, 1% BSA, and 4% SVF. Primary antibodies were incubated for 2 days in blocking buffer at 4°C, and secondary antibodies one day at 4°C. Sections were mounted with homemade Mowiol mounting medium. Antibodies are listed in Supplementary Table 2.

#### Adult brain sections

Floating sections were permeabilized for 3 h at 37°C in a solution of 20% DMSO (Sigma-Aldrich, 276855), 5.75% glycine (Sigma-Aldrich, G7126), and 0.2% Triton X-100 (Sigma-Aldrich, X100) in PBS. Sections were then blocked overnight at 4°C in blocking solution (0.2% gelatin (Sigma-Aldrich, G1393), 0.2% Triton X-100, 0.01% sodium azide (Sigma-Aldrich, S2002) in PBS). Primary antibodies were incubated for 3 days at 4°C, followed by secondary antibodies for 2 days at 4°C in blocking solution. Sections were mounted with homemade Mowiol mounting medium. Antibodies are listed in Supplementary Table 2.

#### Imaging and quantification

Fluorescence images were acquired using a Leica TCS SP8 laser scanning confocal microscope, a Leica DMi8 inverted widefield microscope, or a BioTek Lionheart FX automated microscopy system. Acquisition settings were optimized to avoid spectral overlap between channels. Quantifications were performed using ImageJ/Fiji, AngioTool (providing vessel area, vessel length, and number of branch points), or appropriate software depending on the experiment.

### In situ hybridization (RNAscope)

In situ hybridization was performed on 30 µm PFA-fixed floating brain sections using the RNAscope Multiplex Fluorescent v2 Assay for fixed-frozen tissue samples (Advanced Cell Diagnostics), according to the manufacturer’s instructions. Briefly, floating sections were washed twice in PBS, mounted on Superfrost Plus microscope slides (Fisher Scientific, 12-550-15), and air-dried for 5–10 min at RT. Slides were baked at 60°C for 30 min and fixed in 4% PFA for 15 min. Tissues were dehydrated through a graded ethanol series (50%, 70%, 100%), air-dried, and treated with hydrogen peroxide for 10 min, followed by Protease IV at RT for 30 min.

Probes (Supplementary Table 2) were hybridized at 40°C for 2 h, followed by sequential amplification with RNAscope Multiplex FL v2 Amp 1 (30 min, 40°C), Amp 2 (30 min, 40°C), and Amp 3 (15 min, 40°C). Signal detection was achieved using the appropriate HRP-C1, -C2, or -C3 reagent (15 min, 40°C) followed by the corresponding Opal fluorophore. RNAscope Multiplex FL v2 HRP Blocker was applied for 15 min at 40°C between channels. Slides were subsequently immunostained, mounted and imaged as previously described.

### Single-cell RNA sequencing

Single-cell RNA sequencing (scRNA-seq) experiments were performed on three tissue types, postnatal retinas, postnatal brains, and post-stroke adult brains, each using distinct cell isolation strategies and different library preparation for post-natal and adults’ experiments, as detailed below.

### Postnatal endothelial cells

#### Brain cell isolation and FACS

Postnatal day 10 (P10) brains from five *Sox9iECKO* and *Sox9l/l* mice were collected in ice-cold HBSS without calcium and magnesium. Brains were minced and enzymatically digested using the Neural Tissue Dissociation Kit P (Miltenyi Biotec, 130-092-628): tissues were incubated in Mix 1 for 15 min at 37°C, followed by addition of Mix 2 for an additional 10 min at 37°C. Digestion was stopped with DMEM containing 5% FBS.

Cell suspensions were filtered through a 40 µm cell strainer and centrifuged at 400 × g for 8 min at 4°C. Myelin and cell debris were removed using Debris Removal Solution (Miltenyi Biotec, 130-109-398) and red blood cells were lysed with Red Blood Cell Lysis Solution (Miltenyi Biotec, 130-094-183). After washing in FACS buffer (PBS, 1% FBS, 0.2% EDTA), cells were incubated for 10 min at RT with Fc Block (5 µg/mL CD16/CD32 in FACS buffer; eBioscience, 16-0161-82). Cells were stained for 30 min at RT with APC-conjugated anti-mouse CD31 (BD Biosciences, 551262; 1:200) and FITC-conjugated anti-mouse CD45 (BioLegend, 103108; 1:250) in Fc block solution, then washed and incubated for 30 min at RT with Live/Dead Fixable Aqua (ThermoFisher Scientific, L34957). Compensation controls included unstained cells, single-color controls, FMO controls, and isotype controls (APC Rat IgG2a, κ; BD Biosciences, 551139). After removing dead cells and doublets, CD45− CD31+ endothelial cells were sorted using a BD FACS Aria Fusion.

Sorted cells were then fixed for 24 h at 4°C according to the Chromium Next GEM Single Cell Fixed RNA Sample Preparation Kit protocol (10x Genomics). After fixation and washing, cells were resuspended in PBS containing 0.04% ultra-pure BSA and stored at −80°C until library preparation.

#### Retina cell isolation and enrichment

Retinas from five P10 *Sox9iECKO* and *Sox9l/l* mice were dissected immediately in ice-cold PBS. Retinas were transferred to a pre-heated (37°C) enzyme mix 1 prepared from the Neural Tissue Dissociation Kit P (Miltenyi Biotec, 130-092-628) and digested for 5 min at 37°C. Digestion was stopped by addition of 10% FBS in PBS, and the suspension was filtered through a 35 µm cell strainer (Falcon, 352235). Samples were centrifuged at 300 × g for 3 min at 4°C and the pellet was resuspended in 200 µL PEB buffer (PBS, 0.5 M EDTA, 10% BSA) containing 20 µL anti-mouse CD31 magnetic MicroBeads (Miltenyi Biotec, 130-097-418) and incubated for 20 min at 4°C. After a wash with PEB buffer, endothelial cells were enriched using an MS column (Miltenyi Biotec, 130-042-201) on a MACS MultiStand holder (Miltenyi Biotec, 130-042-303), according to the manufacturer’s instructions.

Enriched cells were then fixed for 24 h at 4°C according to the Chromium Next GEM Single Cell Fixed RNA Sample Preparation Kit protocol (10x Genomics). After fixation and washing, cells were resuspended in PBS containing 0.04% ultra-pure BSA and stored at −80°C until library preparation.

#### Library preparation and sequencing

FLEX libraries were prepared from fixed, sorted brain endothelial cells using the Chromium Next GEM Single Cell Fixed RNA Kit, Mouse Transcriptome (10x Genomics, #1000496) with a target recovery of 10,000 cells per sample. Generated libraries were sequenced on an Illumina NovaSeq6000 S4, followed by demultiplexing and mapping to the mouse transcriptome probe set (Chromium_Mouse_Transcriptome_Probe_Set_v1.1.1_GRCm39-2024-A) using CellRanger multi (pipeline version 9.0.1, 10x Genomics).

#### Bioinformatic analysis

Filtered gene-expression matrices from each sample were processed individually in R (v4.4.0) using Seurat v5.1.0. Seurat objects were created retaining genes in ≥ 3 cells and cells with ≥ 50 features. Data were normalized using SCTransform (regressing on percent_mt and nCount_RNA) and dimensionality reduction was performed using PCA and UMAP (30 PCs). Doublets were identified and removed using DoubletFinder (7.5% doublet rate).

Singlet-filtered datasets were merged and re-normalized. EC were identified based on Cldn5 and Pecam1 pan-EC markers and subclustering was performed after re-normalization, UMAP on 30 PCs, and clustering (resolution = 0.7). EC subtypes were annotated based on established vascular marker genes. Differential gene expression between *Sox9* KO and Ctrl conditions within each EC subtype was assessed using FindMarkers.

### Post-stroke brain endothelial and perivascular cells

#### Cell isolation and FACS

Post-stroke brain tissue was collected from two to three adult mice at 1, 3, or 14 days following tMCAO reperfusion. Each brain was hemisected along the midline to separate the ipsilateral (ischemic) from the contralateral hemisphere. Hemispheres were individually processed using the Neural Tissue Dissociation Kit P (Miltenyi Biotec, 130-092-628) for 15 min in Mix 1 and 10 min in Mix 2 at 37°C. After addition of 5 mL of DMEM/5% FBS to stop the reaction, samples were filtered through a 70 µm strainer. Myelin and debris were removed using Debris Removal Solution (Miltenyi Biotec, 130-109-398) and red blood cells were lysed (Miltenyi Biotec, 130-094-183).

Cells were stained for 30 min at RT with APC-conjugated anti-mouse CD31 (BD Biosciences, 551262), FITC-conjugated anti-mouse CD45 (BioLegend, 103108), and PE-conjugated anti-mouse CD140b/PDGFRβ (BioLegend, 136006) in Fc block solution (1% FBS, 0.2% EDTA, 5 µg/mL CD16/CD32; eBioscience, 16-0161-82 in PBS). Viability was assessed using Live/Dead Fixable Aqua (ThermoFisher Scientific, L34957). After exclusion of dead cells and doublets, viable CD45− CD31+ endothelial cells and CD140b+ perivascular cells were sorted by FACS (BD FACS Aria Fusion).

#### Library preparation and sequencing

Fresh, sorted cell suspensions were resuspended in PBS containing 0.04% ultra-pure BSA. Single-cell libraries were generated using the Chromium Single Cell 3’ Reagent Kit v3.1 (10x Genomics) according to the manufacturer’s instructions, with a target cell recovery of 10,000 cells per library. Generated libraries were sequenced on an Illumina NovaSeq6000 S4, followed by demultiplexing and mapping to the mouse genome (mm10-3.0.0) using CellRanger (pipeline version 3.1.0, 10x Genomics).

#### Bioinformatic analysis

Filtered gene-expression matrices were processed using CellRanger and analyzed in R (v4.4.0) using Seurat v5.1.0. Quality control filtering was applied as follows: cells were retained with nFeature_RNA > 200, nCount_RNA between 1,500 and 25,000, and percent mitochondrial reads < 15%. Cell cycle scoring was performed using the CellCycleScoring function with canonical S-and G2M-phase gene sets. Data were normalized using SCTransform (regressing on percent_mt and nCount_RNA), and dimensionality reduction was performed using PCA and UMAP (25 PCs). Clustering was performed using FindNeighbors (dims = 1:2, reduction = “umap”) and FindClusters (resolution = 0.4). Doublets were identified and removed using DoubletFinder (7.5% doublet formation rate; pN = 0.25, pK = 0.09, PCs = 1:20). Cell type annotation was performed based on expression of canonical marker genes (endothelial cells: Cldn5, Pecam1; mural cells: Kcnj8, Rgs5; astrocytes, fibroblasts, immune cells). Differential gene expression between ipsilateral and contralateral hemispheres, and between control and *Sox9*-overexpressing conditions, was assessed using FindMarkers and FindAllMarkers (Wilcoxon rank-sum test, Bonferroni correction).

### Spatial transcriptomics

#### Sample preparation and library preparation

Spatial transcriptomics was performed on brains from adult control and post-stroke mice (1 day of reperfusion following tMCAO). Brains were embedded in OCT and flash-frozen for cryosectioning at 10 µm thickness. Sections were processed using the Visium Spatial Gene Expression (GEX) Slide & Reagent Kits (10x Genomics, catalog numbers 1000193 and 1000187) according to the manufacturer’s standard protocol.

Tissue sections were placed on GEX Visium slides and stored at −80°C until processing. GEX slides were fixed in methanol and stained with hematoxylin and eosin (H&E) for tissue morphology visualization using an inverted wide DMi8 microscope (Leica). GEX slides were permeabilized for 30 min and incubated with reverse transcription reagents, and cDNA was amplified. Library construction followed the manufacturer’s protocol and included cDNA fragmentation, end repair and A-tailing, adaptor ligation, and sample indexing and amplification. Library quality was assessed by Bioanalyzer (Agilent). Sequencing was performed on an Illumina NovaSeq S4 by Genome Québec.

#### Data processing

Raw sequencing reads were aligned to the mouse reference genome mm10-2020-A using SpaceRanger (v2.1.0, 10x Genomics). Four tissue sections were processed: two control brains (Control_1: 3,167 spots; Control_2: 3,166 spots) and two post-stroke brains at 1 day of reperfusion (Stroke_1: 2,862 spots; Stroke_3: 3,074 spots).

Further processing and analysis were performed in R (v4.4.0) using Seurat v5.1.0. Each section was independently normalized using SCTransform with 3,000 variable features, and dimensionality reduction was performed using PCA and UMAP. Individual Seurat objects were then integrated using Seurat’s integration workflow, producing a merged object comprising 12,269 spots across all four sections with the integrated assay as active assay To identify spatially distinct transcriptional programs, Non-negative Matrix Factorization (NMF) was applied to the integrated dataset at K = 20, using the 3,000 variable features from the integrated assay. Spatial zones were defined based on histological features and NMF program enrichment: control spots were classified as Healthy_Control (Control_1: 3,167; Control_2: 3,166), while stroke spots were classified into Contralateral (Stroke_1: 1,757; Stroke_3: 1,961), Penumbra (Stroke_1: 352; Stroke_3: 389), and Ischemic_Core (Stroke_1: 753; Stroke_3: 724) zones. Differential expression analyses between spatial zones were carried out using FindMarkers (Wilcoxon rank-sum test with Bonferroni correction).

### Quantitative RT-PCR

For isolation of brain endothelial cells, mouse brains were minced and digested for 1 h at 37°C in collagenase/dispase solution (Sigma, 11097113001) supplemented with DNase I (50 µg/mL, DN25) and TCLK (0.147 µg/mL, T7254). Samples were filtered through a 40 µm cell strainer (BD Biosciences) and centrifuged at 400 × g for 10 min at 4°C. Cell pellets were resuspended in 5 mL PBS/25% BSA and centrifuged at 1,000 × g for 20 min at 4°C to remove myelin and debris. After one wash in PBS, cells were incubated with CD31-pre-coated Dynabeads (CD31 antibody: BD Pharmingen, 550274; Dynabeads: ThermoFisher Scientific, 11035) for 15 min at RT. After three magnetic washes, total RNA was extracted using the RNeasy Mini Kit (Qiagen, 74134).

RNA was reverse-transcribed using iScript Reverse Transcription Supermix (Bio-Rad, 1708841). Quantitative PCR was performed using iQ SYBR Green Supermix (Bio-Rad, 1708882). Primers are listed in Supplementary Table 2. Gene expression was normalized to housekeeping genes and relative mRNA levels were calculated using the ΔΔCt method.

### Cell culture and in vitro experiments

#### HUVEC culture and lentiviral transduction

Human Umbilical Vein Endothelial Cells (HUVECs; PromoCell, Cedarlane, C-12203) were cultured in EBM-2 basal medium (Lonza, CC-3121) supplemented with EGM-2 SingleQuots (Lonza, CC-4133). For stable overexpression of *Sox9*, HUVECs were transduced with tTA- or *Sox9*-expressing lentivirus twice at 12 h intervals. Forty-eight hours after the last transduction round, cells were selected with blasticidin (5 µg/mL) or puromycin (1 µg/mL) for 2–3 days. Induction efficiency was confirmed by immunoblotting for SOX9.

#### Proliferation assay

HUVEC proliferation was assessed using the NucBlue Live Cell Stain (ThermoFisher Scientific). Control and *Sox9*-overexpressing cells were thawed and cultured before seeding into 96-well plates. NucBlue was added to each well and incubated for 15–20 min at 37°C. Nuclear counts were acquired daily from day 1 to day 6 using a BioTek Lionheart FX automated microscope with a 4× objective.

#### Invasion assay

Cell invasion was assessed using Corning BioCoat Matrigel Invasion Chambers (Corning, 354578; 8 µm pore diameter, 24-well format). Matrigel matrix (Corning, 354234) was diluted to 200–300 µg/mL in coating buffer (0.01 M Tris pH 8.0, 0.7% NaCl) and applied to insert membranes (0.1 mL per insert) on ice. Coated inserts were incubated at 37°C for 2 h prior to cell seeding. HUVECs were seeded at 2.5 × 104 cells per insert in EBM-2. Complete EGM-2 medium was added to the lower chamber as chemoattractant. After 24 h incubation at 37°C in 5% CO2, non-invaded cells on the apical surface were removed with a moistened cotton swab. Invaded cells on the basolateral membrane were fixed and stained with DAPI and imaged using LX Lionheart automated microscope. Cells were counted and results are expressed as percent invasion through the Matrigel-coated membrane relative to migration through uncoated control inserts.

#### Migration assay

Cell migration was assessed following the invasion protocol above without Matrigel coating. HUVECs were seeded at 2.5 × 104 cells per insert in EBM-2. Complete EGM-2 medium was added to the lower chamber as chemoattractant. After 24 h incubation at 37°C in 5% CO2, non-migrated cells on the apical surface were removed with a moistened cotton swab. Migrated cells on the basolateral membrane were fixed and stained with DAPI and imaged using LX Lionheart automated microscope. Cells were counted and results are expressed as fold change of migrated cells relative to the total number of cells plated.

#### Sprouting assay

HUVEC sprouting was assessed using a fibrin bead assay. HUVECs were trypsinized and mixed with collagen-coated Cytodex microcarrier beads (ratio 2,500 beads per 106 cells; 50 µL bead solution per condition) in warm EGM-2 medium. The mixture was incubated at 37°C for 4 h with agitation every 20 min to allow uniform coating. Bead-coated cells were then transferred to a 6-well plate in 2 mL EGM-2 and incubated at 37°C overnight.

The next day, beads were embedded in a fibrin gel: approximately 15–30 µL of bead suspension (∼100–200 beads per well) was mixed with 45 µL of 10 U/mL thrombin per well, followed by addition of 300 µL fibrinogen solution (3.0 mg/mL in PBS + 100 µg/mL aprotinin) in a 24-well plate. After 15 min at RT and 1 h at 37°C to form the clot, human lung fibroblasts (20,000 cells/well, BJ CRL-2522) were seeded on top of the gel in EGM-2. Medium was changed every other day with complete EGM-2. Plates were fixed and imaged on day 7 using widefield inverted DMi8 microscope. Sprouting was quantified as number of sprouts per bead (approximately 25 beads per condition per experiment).

#### Western blot

Cells were lysed in 1× Laemmli buffer (Sigma-Aldrich, S3401) and denatured at 95°C for 10 min. Protein lysates were separated by SDS-PAGE using precast NP midi gels (ThermoFisher, WG1402BOX) and transferred to nitrocellulose membranes (Bio-Rad, 1620112). Membranes were blocked for 1 h at RT in 5% BSA in TBST, then incubated with primary antibodies overnight at 4°C. After TBST washes, membranes were incubated for 2 h at RT with HRP-conjugated secondary antibodies (Supplementary Table 2). Signals were detected by chemiluminescence using Clarity Western ECL Substrate (Bio-Rad, 1705060S) on a ChemiDoc MP Imaging System (Bio-Rad).

#### Bulk RNA sequencing

Total RNA was extracted from HUVECs (3 control and 3 *Sox9*-overexpressing samples) using RNeasy Mini Plus Kit (Qiagen). RNA quality was assessed using Bioanalyzer. Strand-specific, poly(A)-enriched RNA libraries were prepared using the NEBNext Dual kit (New England Biolabs) and sequenced on an Illumina platform by Genome Québec (project X0124; sequenced 2024-11-15). Differential gene expression analysis was performed using the Biojupies software (Torre et al., Cell Syst 2018).

### Statistical analysis

Statistical analyses were performed using GraphPad Prism 10. All data are presented as mean ± standard error of the mean (SEM). Normality was assessed prior to applying parametric or non-parametric tests. For comparisons between two groups, an unpaired t-test, Welch’s t-test (unequal variances), or Mann-Whitney test was used as appropriate. For multiple group comparisons, one-way ANOVA was used. A p-value < 0.05 was considered statistically significant. For single-cell and spatial transcriptomics, differential expression analyses were carried out using a Wilcoxon rank-sum test with Bonferroni correction for multiple comparisons (as implemented by default in the FindMarkers and FindAllMarkers functions of Seurat). Gene Set Variation Analysis (GSVA) was performed to assess pathway-level activity across conditions; statistical significance of GSVA scores between groups was assessed limma package (Ritchie et al., 2015). A linear model was fit to the GSVA enrichment score matrix using a design matrix encoding experimental groups. Contrast coefficients comparisons were estimated with contrasts.fit, and moderated t-statistics were computed using empirical Bayes variance shrinkage (eBayes). P-values were adjusted for multiple testing using the Benjamini–Hochberg method (FDR correction) via the topTable function.

### Materials availability

Materials are available upon reasonable request.

### Data and code availability

All data that support the findings of this study are available within the article and its supplemental information File and from the corresponding author upon reasonable request. The scRNA-seq, spatial transcriptomics and RNA-seq data discussed herein will be deposited in NCBI’s Gene Expression Omnibus (accession no. xxxx).

## RESULTS

### Brain tip cells show neuro-tip cell characteristics and high *Sox9* transcription factor activity

To define transcriptional regulators of brain endothelial tip cells, we performed single-cell RNA sequencing of endothelial cells (ECs) isolated from P10 mouse brains (**Figure 1a-c**). Initial clustering of FACS-enriched (CD31) brain ECs identified ECs alongside mural, glial, neuronal, and stromal populations, validating successful endothelial enrichment and cell-type annotation (**Supplementary Figure 1a-c**). Sub-clustering and annotation based on established EC markers ^4,7,8^ identified venous (*Tnc*, *Tll1*), proliferative (*Pimreg*, *Troap*), venous capillary (*Lrrc55* and *Rasd2*), capillary (*Cdkn2b* and *Plekhh2*), arterial capillary (*Rhpn2* and *Gask1b*), arterial (*Gja5*, *Gkn3*), and tip cell (*Scn8a* and *Kcnq4*) EC clusters (**Figure 1b, c**). Cell numbers for each cluster are reported in **Data source table**.

**Figure 1:**
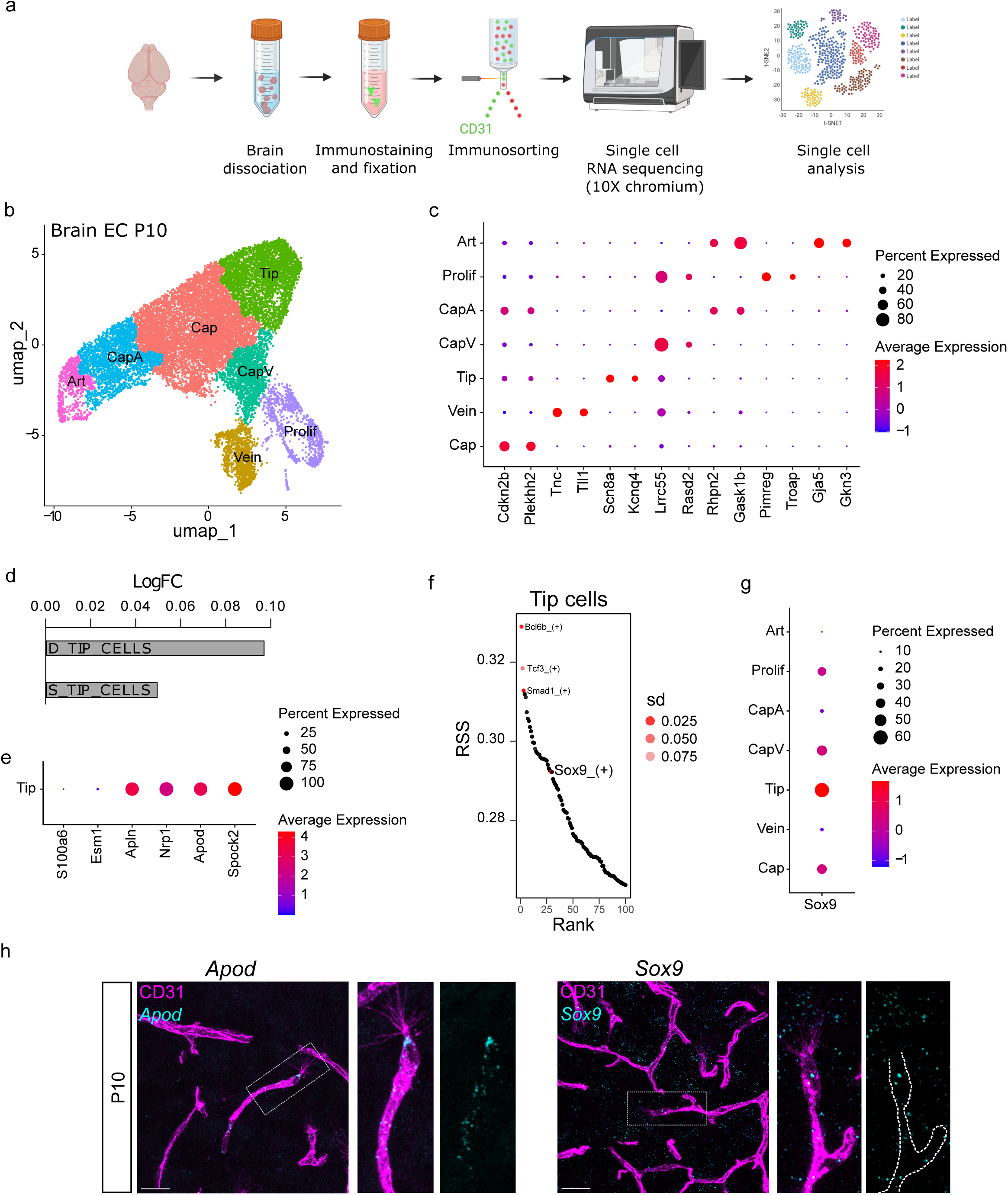
Identification of *Sox9* as a brain tip cell-enriched transcription factor. **a,** Schematic representation of the single-cell experiment using FLEX technology. **b,** UMAP of ECs isolated from P10 mouse brains. **c,** Dot plot showing the two most highly expressed genes in each cluster. **d**, GSVA-based log fold-change scores for gene signatures associated with D-tip or S-tip cell identities in brain endothelial tip cells. **e**, Dot plot showing the expression frequency and level of selected genes in the P10 tip cell cluster. **f**, Transcription factor activity in the P10 tip cell cluster inferred by SCENIC analysis. **g**, Dot plot showing Sox9 expression frequency and level across P10 brain EC clusters. **h**, Representative confocal images of P10 brain tip cells showing *Apod* (left) and *Sox9* (right) mRNA by in situ hybridization combined with CD31 immunostaining. Scale bar 25µm. Art, arterial; CapA, arterial capillary; Cap, capillary; CapV, venous capillary; Prolif, proliferative. Source data are provided as a Source Data file.

Gene set variation analysis (GSVA) revealed that brain tip ECs exhibit a transcriptional profile more closely associated with retinal D-tip rather than S-tip cell identity ^4^, including expression of canonical D-tip markers such as *Apod* and the lack of canonical S-tip markers such as *Esm1* (**Figure 1d, e**). This suggests that brain tip cells acquire a diving identity which parallel their spatial remodelling behavior during post-natal angiogenesis.

To infer upstream regulators of this program, we used SCENIC (Single Cell rEgulatory Network Inference and Clustering) transcription factor activity analysis. Among the top-ranked regulators enriched in tip ECs were *Bcl6b*, *Tcf3*, and *Smad1* (**Figure 1f**). Notably, *Sox9* emerged as one of the most specific transcription factors associated with the tip EC cluster. *Sox9* is a known downstream effector of TGFβ and WNT signaling pathways, which have been implicated in brain endothelial specialization, BBB formation, and neurovascular development ^4,6^. Analysis of transcription factor activity across all endothelial subtypes further revealed that SOX9 regulon activity was largely restricted to tip and capillary ECs, whereas other endothelial populations displayed distinct subtype-specific transcriptional regulators, including strong FOXC2 activity in arterial ECs and TCF-family transcription factors in capillary ECs, suggesting active Notch and WNT signaling in these populations, respectively (**Supplementary Figure 1d**).

Consistent with this, *Sox9* expression was strongly enriched in tip ECs, compared to every other ECs cluster including capillary ECs (**Figure 1g**). In situ hybridization validated both D-tip marker *Apod* and *Sox9* transcripts in P10 brain CD31⁺ tip cells with a correlative expression to single cell data (**Figure 1h**). In accordance with the literature and with our scRNA-seq data (**Supplementary Figure 1e**), *Sox9* was also found in surrounding astrocytes (**Figure 1h**).

To determine whether *Sox9* expression is associated with early acquisition of brain endothelial specialization during the embryonic angiogenic phase, we interrogated a recently published developmental endothelial atlas spanning mouse embryogenesis ^25^. This analysis revealed that *Sox9* expression is selectively enriched in embryonic CNS ECs compared to all other ECs and increases from E10 onwards, paralleling the induction of the brain tip-cell marker *Apod* whereas expression of the S-tip marker *Esm1* decreased over the same developmental window (**Supplementary Figure 1f**). Notably, this temporal increase coincided with the angiogenic process and progressive acquisition of BBB-related transcriptional programs in brain ECs (**Supplementary Figure 1g**), suggesting a developmental transition toward a brain-specialized tip cell state.

Together, these data identify SOX9 as a transcription factor enriched in brain tip cells and associated with their specialized molecular identity.

### Endothelial *Sox9* is required for brain vascularization and brain-tip cell identity

To determine the functional role of *Sox9* in brain ECs, we generated an inducible endothelial-specific knockout model (*Sox9iECKO*) using Apnlr-CreERT2 mice ^23^ (**Supplementary Figure 2a**), which aligned with the venous origin of sprout-forming endothelial cells ^26–28^.

To target the major postnatal window of brain angiogenic sprouting and vascular expansion, tamoxifen was administered at P1, P2, and P3 (**Supplementary Figure 2b**), and mice were analyzed at P10, a stage corresponding to peak postnatal cerebral angiogenesis and active tip cell-mediated vascular remodeling. Efficient deletion of *Sox9* in brain ECs was confirmed by RT-qPCR (**Figure 2a**).

**Figure 2:**
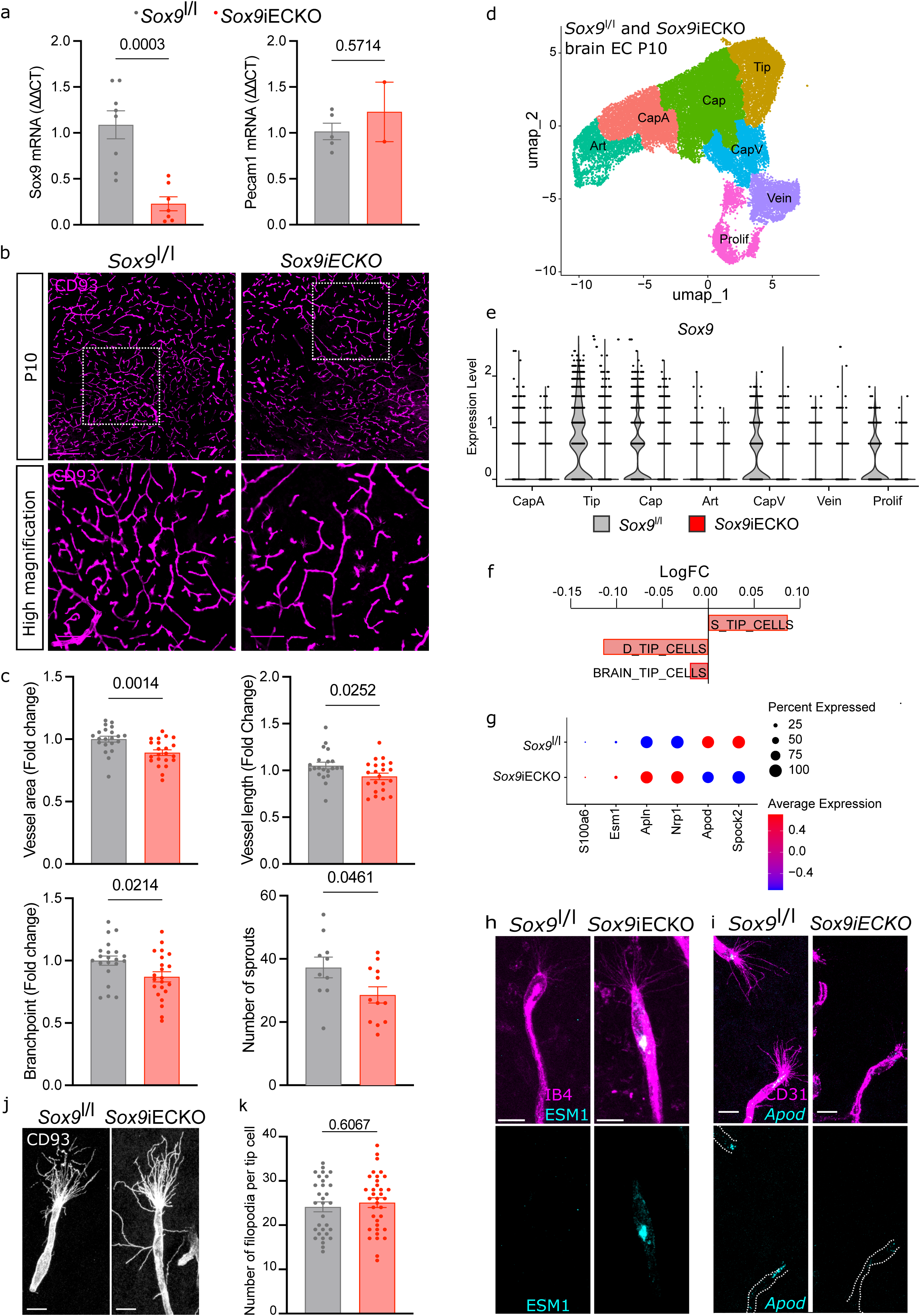
Endothelial *Sox9* is required for brain vascularization and tip cell identity. **a,** RT-qPCR analysis of *Sox9* (left) and *Pecam1* (right) mRNA expression in brain ECs from *Sox9l/l* (grey) and *Sox9iECKO* (red) mice. Sox9, n = 7-8, unpaired t-test with Welch’s correction; Pecam1, n = 2-5, Mann-Whitney test. **b,** CD93 immunostaining of P10 brain coronal sections from *Sox9l/l* (left) and *Sox9iECKO* (right) mice. Scale bars, 200 μm (Top) and 100 μm (Bottom). **c,** Quantification of vessel area, length, branchpoints, and number of angiogenic sprouts in P10 brain coronal sections from *Sox9l/l* (grey) and *Sox9iECKO* (red) mice. n = 10-22; unpaired t-test. **d,** UMAP of P10 brain ECs from *Sox9l/l* and *Sox9iECKO* mice. **e,** Violin plots showing Sox9 expression across EC clusters from *Sox9l/l* (grey) and *Sox9iECKO* (red) mice. p<0,001. **f**, GSVA score of gene set signatures involved in S-, D- and brain-tip cells identity represented by LogFC fold change from *Sox9*iECKO compared to *Sox9*^l/l^ tip EC. p<0,001. **g,** Dot plot showing the expression of selected genes in *Sox9l/l* and *Sox9iECKO* tip cell clusters. **h-i,** Representative images of P10 brain tip cells co-stained for IB4 and ESM1 (I) or CD31 and *Apod* (J) in *Sox9l/l* (left) and *Sox9iECKO* (right) mice. Scale bars, 10 μm. **j,** Representative images of P10 brain tip cells from *Sox9l/l* (left) and *Sox9iECKO* (right) mice immunostained for CD93. Scale bars, 10 μm. **k,** Quantification of the number of filopodia per tip cell in *Sox9l/l* (grey) and *Sox9iECKO* (red) mice following CD93 immunostaining. n = 31-35; Mann-Whitney test. Source data are provided as a Source Data file.

Endothelial deletion of *Sox9* at birth resulted in a significant reduction of vascular network complexity in the thalamic area, characterized by decreased vessel area, vessel length, branchpoints, and number of angiogenic sprouts (**Figure 2b, c**). A similar but milder phenotype was observed in the cortical area (**Supplementary Figure 2c, d**), without changes in body or brain weights (**Supplementary Figure 2e**).

To investigate the cellular basis of this phenotype, we performed scRNA-seq of FACS-isolated brain ECs at P10. All major EC populations were identified, and efficient loss of *Sox9* expression and activity was confirmed (**Figure 2d, e and Supplementary Figure 2f-h**). Pathway analysis revealed a significant decrease in brain- and D-tip cell signatures and a concomitant increase in S-tip cell signatures in *Sox9*-deficient tip ECs (**Figure 2f**).

Accordingly, *Sox9* deletion led to reduced expression of D-tip markers such as *Apod* and induction of S-tip markers, including *Esm1*, which is normally absent from brain tip ECs (**Figure 2g-i**). Notably, the transcriptional impact of SOX9 loss was largely restricted to the tip EC cluster, which exhibited the highest number of differentially expressed genes, whereas other EC populations were minimally affected (**Supplementary Figure 2i**).

Despite these transcriptional and vascular defects, *Sox9* deletion did not alter filopodia formation in tip cells (**Figure 2j, k**), suggesting that SOX9 primarily regulates tip-cell identity and function rather than morphological features of sprouting.

### SOX9 controls neuroretina vascularization and D-tip cell identity

Given the shared mechanisms of CNS angiogenesis, we next examined the role of SOX9 in retinal vascular development. Using a previously published dataset of scRNA-seq from P6 and P10 retina ^4^, we first confirmed that *Sox9* ranks among the top transcription factors active in D-tip but not S-tip retinal ECs, correlated with a high and selective expression to the D-tip cell cluster (**Supplementary Figure 3a, b**).

Endothelial *Sox9* deletion at birth resulted in a marked reduction in vascular density and complexity of the deep retinal plexus at P10, including decreased vessel area, length, branchpoints, and D-tip cell number (**Figure 3a, b and Supplementary Figure 3c**), without affecting superficial plexus development at P6 (**Supplementary Figure 3d, e**).

**Figure 3:**
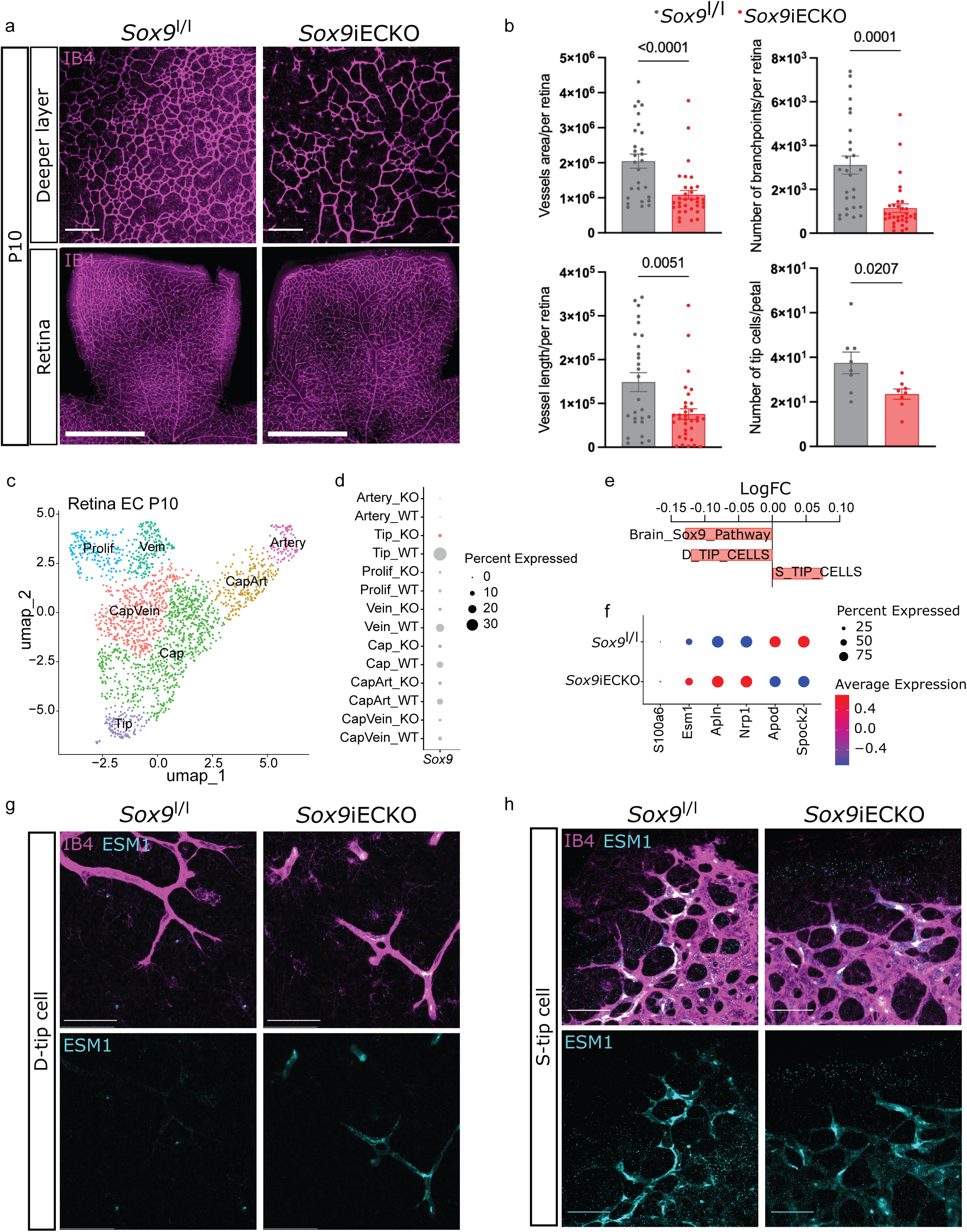
Endothelial *Sox9* regulates neuro-retina vascularization and D-tip cell identity. **a,** IB4 immunostaining of P10 whole-mount retinas from *Sox9l/l* (left) and *Sox9iECKO* (right) mice. Top panels show the deep vascular layer only. Scale bars, 150 μm (Top) and 500 μm (Bottom). **b,** Quantification of vessel area, length, branchpoints, and number of angiogenic sprouts in the deep vascular layer of P10 retinas from *Sox9l/l* (grey) and *Sox9iECKO* (red) mice. n = 27-33; Mann-Whitney test (Vessel area), or unpaired t-test with (Branchpoint, Vessel length) or without Welch’s correction (Number of tip cells). **c**, UMAP of P10 retina EC of *Sox9*^l/l^ and *Sox9*iECKO single cell data. **d,** Dot plot showing *Sox9* expression across EC clusters from *Sox9l/l* (grey) and *Sox9iECKO* (red) retinas. **e,** GSVA scores for the brain *Sox9*-dependent-, S- and D-tip cell gene signatures in *Sox9iECKO* versus *Sox9l/l* retinal tip endothelial cells. **f,** Dot plot showing the expression of selected genes in *Sox9l/l* and *Sox9iECKO* tip cell clusters. **g-h,** Representative images of P10 retinal D-tip cells (G) or S-tip cells (H) co-stained for IB4 and ESM1 in *Sox9l/l* (left) and *Sox9iECKO* (right) mice. Scale bars, 50 μm. Source data are provided as a Source Data file.

scRNA-seq of retinal *Sox9*^l/l^ and *Sox9*iECKO ECs confirmed that *Sox9* deletion induced a transcriptional shift with reduced D-tip cell signature and brain-generated *Sox9* pathway and increased S-tip cell signature (**Figure 3c-e and Supplementary Figure 3f**). This was accompanied by decreased expression of D-tip markers and increased expression of S-tip markers such as *Esm1* (**Figure 3f-h**).

These findings show that SOX9 regulates D-tip cell specification and neuroretina vascularization.

### SOX9 promotes *in vivo* and *in vitro* endothelial sprouting and matrix remodeling programs

To define SOX9-dependent pathways, we analyzed gene signatures in *Sox9*-deficient ECs. In the brain, loss of *Sox9* resulted in reduced activity of various pathways associated with sprouting angiogenesis, and neuro-tip cell characteristics such as extracellular matrix remodeling, and BBB-related functions (**Figure 4a, c**). Overall, a similar effect was observed in the retina, except for the extracellular matrix pathways that were increased after *Sox9* deletion (**Figure 4b, d**).

**Figure 4:**
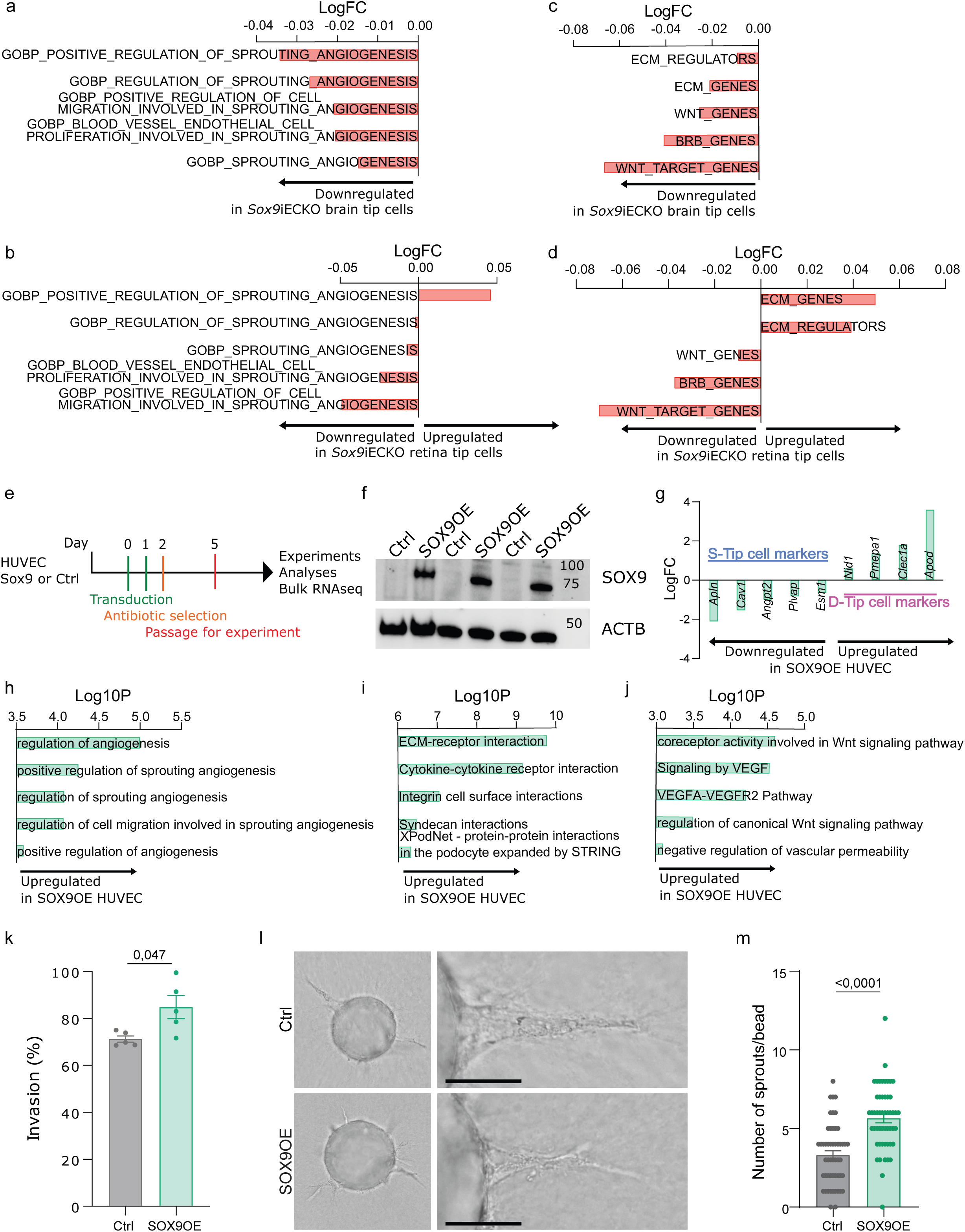
*Sox9* regulates endothelial sprouting *in vivo* and *in vitro*. **A-b,** GSVA scores of selected gene set signatures related to sprouting angiogenesis in *Sox9*iECKO compared to *Sox9*^l/l^ brain (a) and retina (b) tip ECs. p<0,001 for brain data. Retinas, not significant. **c-d,** GSVA scores for D-tip cell-related gene set signatures in *Sox9*iECKO compared to *Sox9*^l/l^ brain (c) and retina (d) tip ECs. p<0,001 for brain data. Retinas, not significant. **e,** Schematic of the experimental timeline in control and SOX9-overexpressing (SOX9OE) HUVECs. **f,** Immunoblot analysis of SOX9 and ACTB in control and SOX9OE HUVECs. **g,** Histogram showing selected differentially expressed genes in SOX9OE HUVECs relative to control cells. p<0,001. **h-j,** GSVA scores for selected gene signatures related to sprouting angiogenesis (h), cell interaction (i), and permeability (j) in SOX9OE versus control HUVECs. p<0,005. **k,** Percentage of invasion of control and SOX9OE HUVECs. n=5, unpaired t-test with Welch’s correction. **l-m**, Representative images (l) and quantification (m) of control and SOX9OE HUVECs *in vitro* sprouting assay. n=50, N=2, Mann-Whtiney test. Scale bar, 50 μm. Source data are provided as a Source Data file.

To test whether SOX9 is sufficient to induce this state and because primary brain ECs rapidly lose their specialized phenotype *in vitro* ^29^, we overexpressed *Sox9* (SOX9OE) in human umbilical vein endothelial cells (HUVECs) as they display transcriptional features resembling S-tip cells in EGM-2 medium (**Figure 4e**).

After validation of the expression by immunoblotting (**Figure 4f**), transcriptomic analysis revealed that SOX9OE induces a robust endothelial reprogramming (**Supplementary Figure 4a)**, characterized by increased expression of neuro-tip markers such as *Apod,* and suppression of S-tip markers, including *Esm1* (**Figure 4g**,). Pathway analysis showed increased activity of sprouting angiogenesis, matrix remodeling, WNT, and VEGF signaling pathways (**Figure 4h-j**).

Functionally, SOX9 overexpression enhanced endothelial invasive capacity in transwell invasion assays (**Figure 4k**), migration (**Supplementary Figure 4b, c**), and enhanced sprouting 3D *in vitro* microcarrier bead sprouting assays (**Figure 4l, m**), without affecting proliferation (**Supplementary Figure 4d**). These data indicate that overexpressing *Sox9 in vitro* promotes an angiogenic and matrix-remodeling endothelial state associated with neuro-tip cell identity.

### Post-stroke angiogenic ECs lack SOX9-dependent neuro-tip cell identity

To investigate whether SOX9-dependent programs can be reactivated *in vivo* during pathological angiogenesis, we performed scRNA-seq of ECs isolated from control, contralateral, and ipsilateral hemispheres following transient middle cerebral artery occlusion (tMCAO) after 1, 3, and 14 days of reperfusion (**Figure 5a, b**).

**Figure 5:**
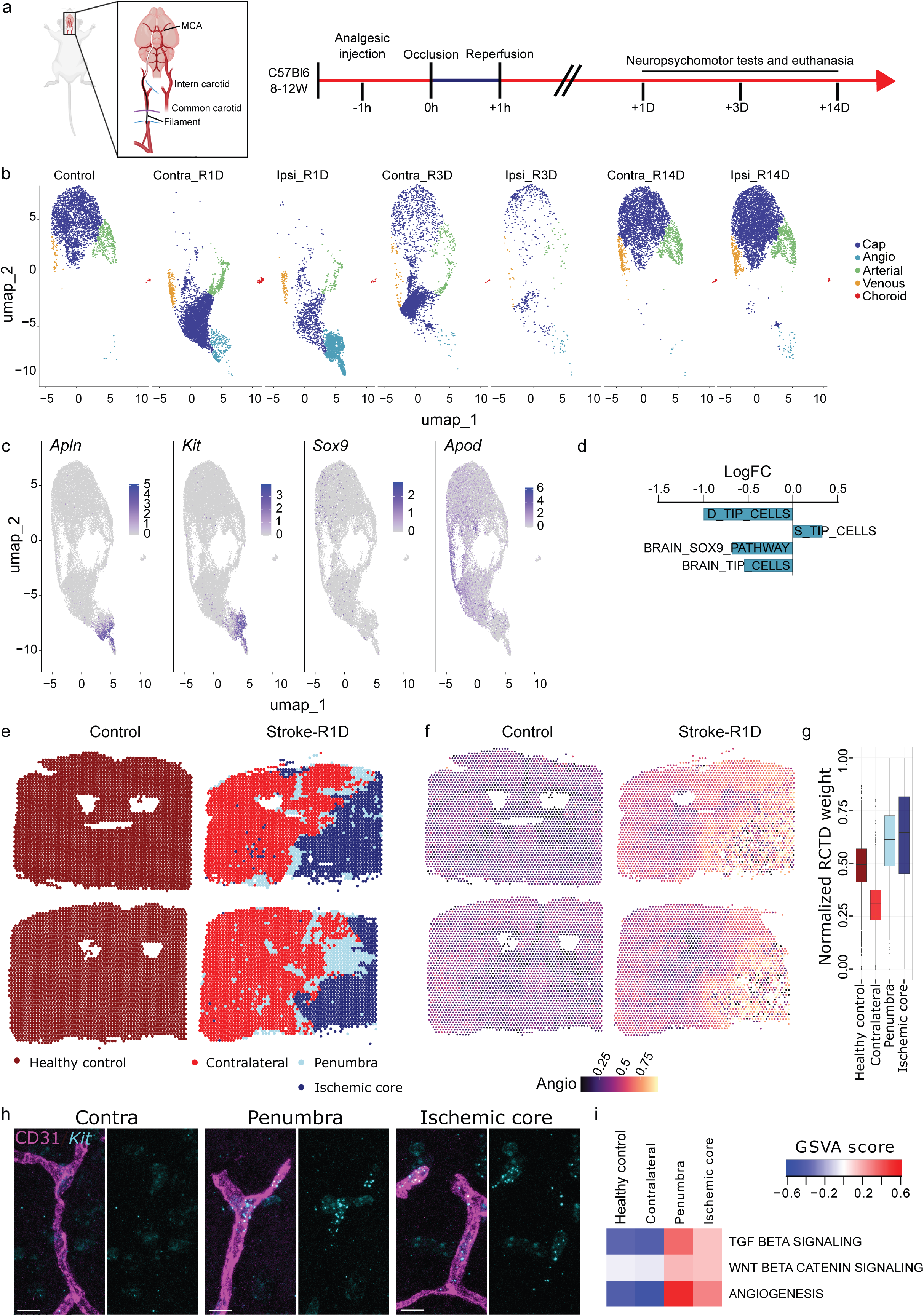
Post-stroke angiogenic ECs lack *Sox9*. **a,** Schematic of the transient middle cerebral artery occlusion (tMCAO) model and experimental timeline in C57BL/6 mice. **b,** UMAP of ECs from control, contralateral (Contra), and ipsilateral (Ipsi) tMCAO brain hemispheres after 1, 3, or 14 days of reperfusion. **c,** Feature plots showing the expression of selected genes. **d,** GSVA scores for the D-, S-tip cells, and brain *Sox9*-dependent-, Brain-tip gene signatures in the angiogenic EC cluster. p<0,001. **e**, Spatial zonation of the different regions after tMCAO. **f-g**, Spatial distribution (f) and normalized RCTD weight (g) representation of the angiogenic EC cluster. **h**, Representative images of contralateral (left), penumbra (middle) and ischemic core (right) regions of a coronal brain section at 1 day of reperfusion stained for CD31 and *Kit* mRNA by in situ hybridization. Scale bars, 10 μm. **i,** GSVA scores for selected gene signatures related to angiogenic signaling and angiogenesis in the stroke zonations. Source data are provided as a Source Data file.

In control adult brains, ECs were organized along arterial, capillary, and venous axes, with no detectable proliferative or tip cell populations, consistent with a quiescent vasculature (**Supplementary Figure 5a, b**). Following stroke, we observed marked transcriptional remodeling of ECs, as both ipsi- and contralateral ECs shifted to a new state 1-day post-reperfusion. These stroke-induced changes progressively resolved 3-days post-reperfusion to return to near baseline by day 14 (**Figure 5b**). Gene set activity analyses of 1-day post-reperfusion compared to control ECs revealed increased hypoxia and vascular permeability signatures across EC populations (capillary, artery and vein) (**Supplementary Figure 5d)**. These data were correlated with an increase of Hif1a hypoxic marker and decrease of Mfsd2a BBB marker expression (**Supplementary Figure 5e**), confirming stroke-induced vascular stress.

Clustering and marker analysis also identify the emergence of a distinct endothelial population enriched in the ipsilateral hemisphere at day 1 post-reperfusion (**Figure 5b, Supplementary Figure 5c**), which we termed angiogenic ECs. This cluster expressed canonical angiogenic markers such as *Apln* and *Kit*, which were largely absent from other EC populations (**Figure 5c and Supplementary Figure 5b**). However, in contrast to post-natal tip cells, these pathological angiogenic ECs lacked expression of *Sox9* and neuro-tip markers such as *Apod* (**Figure 5c, Supplementary Figure 5f**).

Consistent with this, angiogenic ECs displayed reduced D-tip, brain-tip, and SOX9-dependent transcriptional signatures, together with relative enrichment of S-tip-like signatures compared to developmental brain-tip cells (**Figure 5d and Supplementary Figure 5f**). Direct comparison with P10 developmental brain tip ECs further confirmed that post-stroke angiogenic ECs fail to recapitulate the full neuro-tip cell transcriptional program despite angiogenic activation, including a marked reduction in matrix remodeling, WNT signaling, and BBB-associated pathways (**Supplementary Figure 5g, h**). These findings indicate an incomplete and aberrant activation of angiogenic programs in the adult injured brain.

To determine whether this angiogenic endothelial population is spatially restricted to ischemic regions, we next integrated spatial transcriptomic analyses of stroke sections at 1-day post-reperfusion and control (n=2 per condition). Unsupervised non-negative matrix factorization (NMF)-based spatial domain analysis identified transcriptionally distinct regions within the control, contra-, and ipsilateral hemisphere (**Figure 5e**), corresponding to the ischemic core and penumbra, which were further validated by expression of canonical marker genes.

We observed a marked reduction in the number of detected genes (nFeatures) within the ischemic core, and the penumbra, compared to contralateral and control regions, consistent with tissue damage and transcriptional failure (**Supplementary Figure 5i**). Aligning with the spatial organization, pathway activity analysis further validated stroke-associated hallmark pathway enrichment in the penumbra and ischemic core (**Supplementary Figure 5j**).

Deconvolution of single-cell ECs onto spatial transcriptomic data revealed that angiogenic ECs preferentially localize in the penumbra and ischemic core (**Figure 5f, g**). Importantly, ISH analysis of *Kit* expression confirmed the angiogenic ECs localization (**Figure 5h**).

Finally, analysis of angiogenic signaling pathways across stroke zonation demonstrated that the penumbra and ischemic core exhibit robust activation of angiogenic and sprouting-related programs (**Figure 5i**). However, the angiogenic ECs emerging within a pro-angiogenic microenvironment, were lacking SOX9-dependent brain-tip cell identity which was *in fine* correlated with the absence of tip cells after stroke (**Figure 5d**).

Together, these findings demonstrate that post-stroke angiogenic ECs arise specifically within ischemic territories but fail to acquire the specialized SOX9-dependent transcriptional program characteristic of developmental brain-tip cells subsequently failing to induce genuine tip cell specification.

## DISCUSSION

Our study identifies SOX9 as a regulator of organotypic endothelial tip cell specification in the developing CNS. By combining single-cell and genetic loss-of-function and overexpression models *in vivo* and *in vitro*, we show that SOX9 is selectively enriched in neurovascular tip cells and is required for the acquisition of their specialized transcriptional and functional programs. Endothelial deletion of *Sox9* impairs postnatal cerebral and neuroretinal vascularization, disrupts neuro-tip cell transcriptional programs, and reduces angiogenic sprouting and matrix remodeling pathways. Conversely, SOX9 expression is sufficient to promote a neuro-tip-like state associated with enhanced invasion and sprouting in human endothelial cells. Finally, we show that angiogenic endothelial cells emerging one day after ischemic stroke fail to reactivate *Sox9* and acquire the specialized developmental neuro-tip cell specification despite exposure to a pro-angiogenic environment. Together, these findings identify SOX9 as a key transcriptional modulator of the CNS vascularization that could promote vascular regeneration after stroke injury.

Our results extend previous work demonstrating that endothelial tip cells are heterogeneous and acquire tissue-specific molecular identities during vascular development ^4–6^. Classical angiogenic tip cells are primarily defined by VEGF responsiveness and expression of canonical markers such as *Esm1*, *Apln*, and *Dll4* ^1–3^. However, recent work demonstrated that retinal endothelial tip cells invading the neuroretina acquire a distinct D-tip cell identity characterized by activation of TGFβ signaling, expression of blood-retina barrier-associated genes, and repression of canonical superficial tip cell markers such as *Esm1* ^4^. A conceptual advance of this work is the demonstration that CNS tip cells are not simply migratory angiogenic endothelial cells, but a specialized population with a distinct transcriptional state. In the developmental brain, tip cells align more closely with retinal D-tip cells than superficial S-tip cells, including enrichment of *Apod* and suppression of *Esm1*, consistent with a shared neurovascular program across CNS vascular beds. This convergence supports the idea that angiogenesis in neural tissues is guided by a conserved tip-cell identity that is tailored to barrier formation, sprouting, and tissue maturation rather than by generic endothelial expansion alone. Importantly, our data identify SOX9 as a major transcriptional regulator associated with this specialized endothelial program.

SOX9 is classically known for its essential roles in embryonic development and cancer, where it induces cell specification programs in various organs from all three germ layers. Among others: chondrogenesis, testis differentiation, cardiac valve development, neural crest development, stem cell maintenance, and lung development ^30,31^. While SOX9 expression is generally lost after developmental stages, it is described to be re-expressed in adulthood to increase cancer proliferation and metastasis ^31^. In the vasculature, SOX9 has been implicated in endovascular progenitors’ fate, EndMT, and fibrosis formation *in vivo* and *in vitro* ^32–34^. However, its role in CNS endothelial cells remains unexplored. Our data suggest that SOX9 acts downstream of developmental signaling pathways implicated in neurovascular specialization. Prior work established a requirement for TGFβ signaling in retinal D-tip specification and neuroretina vascularization ^4^, while WNT/β-catenin signaling is recognized as a core driver of brain angiogenesis, blood-brain barrier development, and endothelial specialization in the CNS ^30,35,36^. Although our study does not formally place SOX9 within a linear signaling hierarchy, the selective enrichment of SOX9 in developmental neuro-tip cells, together with its ability to drive neuro-tip gene expression, matrix remodeling, and sprouting programs, is consistent with a model in which SOX9 integrates developmental signaling pathways to coordinate angiogenic sprouting with the acquisition of CNS-specific endothelial properties.

Interestingly, *Sox9* deletion primarily affected endothelial transcriptional identity and vascular complexity without significantly altering filopodia formation in the remaining tip cells. This finding aligns with the concept that morphological and molecular tip cell features are distinctively regulated ^37^. Our data suggest that endothelial tip cells first acquire a generic angiogenic state with filopodia formation and subsequently undergo tissue-specific transcriptional specialization. In neural tissues, this specialization appears to involve activation of matrix-remodeling, barrier-associated, and neurovascular sprouting programs under SOX9 control.

The strong regulation of extracellular matrix and invasion-associated pathways by SOX9 further supports this interpretation. Developmental CNS angiogenesis requires endothelial cells to migrate through highly specialized neural microenvironments while simultaneously establishing barrier properties and interactions with perivascular, neural, and glial cells ^5,38–40^. Consistent with this, *Sox9*-deficient endothelial cells displayed reduced extracellular matrix remodeling signatures *in vivo*, whereas SOX9 overexpression promoted endothelial invasion, migration, and sprouting *in vitro*. SOX9 is known in other systems to regulate extracellular matrix-associated genes and tissue remodeling programs, including collagens, matrix metalloproteinases, and adhesion molecules ^30,31^. These features are also hallmarks of the EndMT process which was recently shown to be partially induce by SOX9 in HUVECs ^32^. Our findings, therefore, suggest that SOX9 enables neuro-tip endothelial cells to adapt to the unique extracellular environment of the developing brain and neuroretina. An important aspect of our study is the comparison between developmental angiogenesis and injury-induced vascular remodeling after ischemic stroke, showing their distinction. While numerous reviews highlight the role of post-stroke angiogenesis and its therapeutic potential to improve the life-threatening outcomes of cerebral ischemia, the literature also points to a lack of direct evidence for sprouting angiogenesis and of the molecular characteristics of post-stroke angiogenesis ^19,20,41,42^. Following a stroke, we identify a transient angiogenic endothelial population emerging specifically within the ischemic core and penumbra. Despite activation of angiogenic pathways and a pro-angiogenic environment, aligning with the previous studies ^19,20,41,42^, we show that these cells do not express *Sox9* and fail to acquire the full developmental neuro-tip program. Instead, they retain a partial angiogenic response and features more closely related to non-specialized tip endothelial cells with defective induction of matrix remodeling, WNT-related, and barrier-associated pathways compared to developmental brain tip cells. This suggests that post-stroke angiogenesis recapitulates only part of the developmental angiogenic program and fails to engage the correct CNS endothelial specification program. Developmental context-dependent mechanisms, potentially involving neuronal, glial, or extracellular matrix-derived signals, may be required to activate SOX9 and establish proper neuro-tip identity. These findings may help explain why post-stroke angiogenesis often results in incomplete vascular recovery and persistent blood-brain barrier dysfunction.

Several limitations should be considered. First, although the data support a strong association between TGFβ, SOX9, and neuro-tip identity, the direct molecular mechanism that activates SOX9 *in vivo* remains to be fully defined. Similarly, the transcriptional targets through which SOX9 regulates matrix remodeling and endothelial specialization require further investigation. Future chromatin accessibility and SOX9 occupancy studies will be important to define the regulatory landscape controlled by SOX9 in CNS endothelial cells. Second, while our overexpression experiments show that SOX9 is sufficient to induce key features of the program in HUVECs, they do not establish that SOX9 alone is sufficient to generate a fully faithful neuro-tip cell *in vivo*. Third, the stroke analysis reveals an incomplete angiogenic state, but further work will be needed to determine whether restoring SOX9 activity can improve vascular function, BBB recovery, or tissue repair after ischemic injury.

In conclusion, our study identifies SOX9 as a determinant regulator of organotypic neurovascular tip-cell identity, which is required for proper brain and retinal vascularization. More broadly, our study supports the concept that organ-specific endothelial specialization is actively coordinated during angiogenesis through dedicated transcriptional programs. Post-stroke angiogenic EC’ failure to reactivate SOX9-dependent neuro-tip programs further reveals a fundamental difference between developmental and pathological angiogenesis in the CNS. By linking SOX9 to the acquisition of a specialized sprouting state in the brain and retina, our findings advance the understanding of how endothelial heterogeneity is established during CNS vascularization and suggest that targeting pathways that promote neuro-tip endothelial specialization may therefore represent a promising strategy to improve functional vascular regeneration and neurovascular repair in ischemic and neurovascular diseases.

## Author contributions

TA and AD conceptualized and designed the study. TA conducted histological experiments. SRT and BL produced the virus. TA performed single-cell transcriptomic experiments, TA, JPH, and GC analyzed the bioinformatic data. TA and LK analyzed the data. AD secured funding for the project. TA and AD prepared the initial draft of the manuscript, and all co-authors contributed to reviewing and editing the final text.

## Statements and Declarations

## Acknowledgments

The authors gratefully acknowledge Atik Fuad and Anais Defois for their supportive contributions to this work. Our sincere thanks to the animal facility for their dedicated care of the animals used in this study. This work was supported by project grants from the Canadian Institutes of Health Research (2019PJT-165871, 202203PJT-183658, 202403PJT-517269) and the Natural Sciences and Engineering Research Council of Canada (RGPIN-2022-04726). T.A. was a recipient of fellowships from Fond de Recherche en Ophtalmologie de l’Université de Montreal (FROUM 2022 and 2023) and Fonds de Recherche du Québec en Sante (FRQS 330166 and 375085).

Competing Interests

Authors declare no competing interests for this study.

## Novelty and Significance

What is known?

- Angiogenic endothelial tip cells guide vascular sprouting during development and tissue repair.
- Recent studies have demonstrated that endothelial tip cells acquire organ-specific identities required for proper tissue vascularization.
- The molecular mechanisms linking angiogenic sprouting to brain- and retina-specific endothelial specialization remain poorly understood.

What new information does this article contribute?

- SOX9 is a transcription factor specifically enriched in cerebral and neuroretinal endothelial tip cells.
- Endothelial SOX9 is required for neural tissue vascularization and promotes tip cell programs associated with sprouting angiogenesis, matrix remodeling, and endothelial specialization.
- Post-stroke angiogenic endothelial cells fail to reactivate the SOX9-dependent developmental tip cell program despite robust angiogenic activation.

This study identifies SOX9 as a transcriptional regulator of specialized endothelial tip cell in the developing brain and retina. Using single-cell, spatial, and bulk transcriptomic analyses combined with endothelial-specific genetic loss-of-function and overexpression models, we demonstrate that SOX9 promotes endothelial programs required for neurovascular angiogenesis and tissue-specific vascular specialization. These findings provide a molecular mechanism underlying the concept of organotypic tip cell specification that we and others have recently described in neural tissues. Importantly, we show that angiogenic endothelial cells emerging after ischemic stroke do not reactivate the SOX9-dependent developmental program, revealing fundamental differences between developmental and regenerative angiogenesis. Together, our findings establish SOX9 as a key regulator of neurovascular tip cell identity and suggest that restoring developmental endothelial specification may represent a strategy to improve vascular regeneration after ischemic injury.

